# mLiftOver: Harmonizing Data Across Infinium DNA Methylation Platforms

**DOI:** 10.1101/2024.03.18.585415

**Authors:** Brian H. Chen, Wanding Zhou

## Abstract

Infinium DNA methylation BeadChips are widely used for genome-wide DNA methylation profiling at the population scale. Recent updates to probe content and naming conventions in the EPIC version 2 (EPICv2) arrays have complicated integrating new data with previous Infinium array platforms, such as the EPIC and the HumanMethylation450 (HM450) BeadChip. We present *mLiftOver*, a user-friendly tool that transfers probe ID, methylation level, and signal intensity data across different Infinium platforms. It manages probe replicates, missing data imputation, and platform-specific bias for accurate data conversion. We validated the tool by applying HM450-based cancer classifiers to EPICv2 cancer data, achieving high accuracy. Additionally, we successfully integrated EPICv2 healthy tissue data with legacy HM450 data for tissue identity analysis and produced consistent copy number profiles in cancer cells.

**Availability and implementation:** mLiftOver is implemented R and available in the Bioconductor package SeSAMe (version 3.21.13+): https://bioconductor.org/packages/release/bioc/html/sesame.html Analysis of EPIC and EPICv2 platform-specific bias and high-confidence mapping is available at https://github.com/zhou-lab/InfiniumAnnotationV1/blob/main/Anno/EPICv2/EPICv2ToEPIC_conversion.tsv.gz The source code is available at https://github.com/zwdzwd/sesame/blob/devel/R/mLiftOver.R under the MIT license.

## 1 INTRODUCTION

The Infinium DNA methylation BeadChips (Illumina, Inc., San Diego, CA, USA) are widely used tools for large-scale DNA methylation profiling, contributing significantly to our understanding of DNA methylation biology over the past two decades (Bibikova *et al*., 2006). Combining bisulfite conversion with a genotyping array technology, this platform has been instrumental for consortia projects, such as The Cancer Genome Atlas (TCGA), and has accumulated over 80,000 HM450 samples (Maden *et al*., 2021) and a comparable number of EPIC array methylation profiles in the Gene Expression Omnibus (GEO). Designed for population research, including meQTL studies (Hawe *et al*., 2022; Min *et al*., 2021), epigenetic risk scoring (Thompson *et al*., 2022; Aref□Eshghi *et al*., 2020), and epigenome-wide association studies (EWAS) (Battram *et al*., 2022; Li *et al*., 2019), Infinium arrays offer cost-effectiveness, high quantitative resolution, ease of use, and the ability to accommodate a wide range of DNA inputs (Lee *et al*., 2024). Their high throughput capabilities have accelerated clinical applications in areas such as cancer diagnosis (Capper *et al*., 2018), liquid biopsies (Li *et al*., 2022), and forensic science (Mannens *et al*., 2022). More recently, this technology has supported the creation of an extensive pan-mammalian DNA methylome atlas (Haghani *et al*., 2023; Ding *et al*., 2023; Arneson *et al*., 2022).

The arrays’ probe naming system (*i*.*e*., *cg* number), beginning with the Infinium HumanMethylation27 BeadChip (HM27), has been a cornerstone for cross-referencing probes with unique CpG sites within the genome. Each *cg* number corresponds to a unique 122-mer sequence centered on the target cytosine-guanine dinucleotide (CpG site), with array probes designed against these sequences. Originally, the Infinium arrays featured a one-to-one design—one probe set per 122-mer sequence—enabling a unique mapping to the human genome and facilitating cross-referencing 122-mer IDs, or *cg* numbers, with genomic CpG locations. This method of referencing, common in EWAS literature (Battram *et al*., 2022; Xiong *et al*., 2020) and studies of methylation-genotype interactions (Hawe *et al*., 2022; Min *et al*., 2021), provided a convenient yet imperfect system for indexing probe sequences or CpG sites within a genome assembly.

The main limitation of the original *cg* number system arises from its non-specificity—a single *cg* number could correspond to multiple probe designs targeting the same 122-mer sequence. Additionally, this framework did not allow the inclusion of multiple replicate probes (Bibikova *et al*., 2009), which would enhance the robustness of measurements. With the advent of newer Infinium array generations like the EPICv2 (Noguera□Castells *et al*., 2023; Kaur *et al*., 2023) and other non-human arrays (Zhou *et al*., 2022; Arneson *et al*., 2022), a more precise naming system was introduced. This new system retains the *cg* number as a prefix but adds additional information to distinguish between probes, accounting for Infinium chemistry, strand orientation, and replicate indices (Zhou *et al*., 2022). However, the introduction of additional probe details, while methodologically sound, can impede the integration of newly generated methylation data with legacy datasets using the antiquated probe naming system.

Moreover, the static probe content selection in Infinium technology reflects the evolving understanding of methylation biology (Zhou *et al*., 2017). Each array generation—HM27, HM450, EPIC, and EPICv2—has refined probe content to represent better emerging biological insights, like gene body methylation (Yang *et al*., 2014) and *cis*-regulatory element methylation (Neiman *et al*., 2017). However, integrating legacy data generated on previous platforms with missing probes remains technically challenging, especially for applications like epigenetic clocks (Horvath, 2013) and cancer classification models (Capper *et al*., 2018), which require specific CpGs in a model. Although data imputation strategies can help fill missing values within samples, many methods, such as matrix factorization (Mazumder *et al*., 2010), cannot accommodate the complete missingness of a specific probe in the query dataset. How to continue leveraging the legacy data and predictive models on the ever-evolving Infinium platforms has become a pressing technical need.

To respond to this need, we introduce *methylation LiftOver (mLiftOver)*, a tool designed to harmonize Infinium data efficiently across platforms, including the EPICv2 array. *mLiftOver*, handles ID conversion, replicate probe measurement resolution, and missing data imputation (**Figure 1A**). It is compatible with the R/Bioconductor ecosystem and enables data conversion with varying stringency levels. We demonstrate its utility by applying it to public EPICv2 datasets, showcasing its high performance and utility in bridging different Infinium platforms.

**Figure 1.**
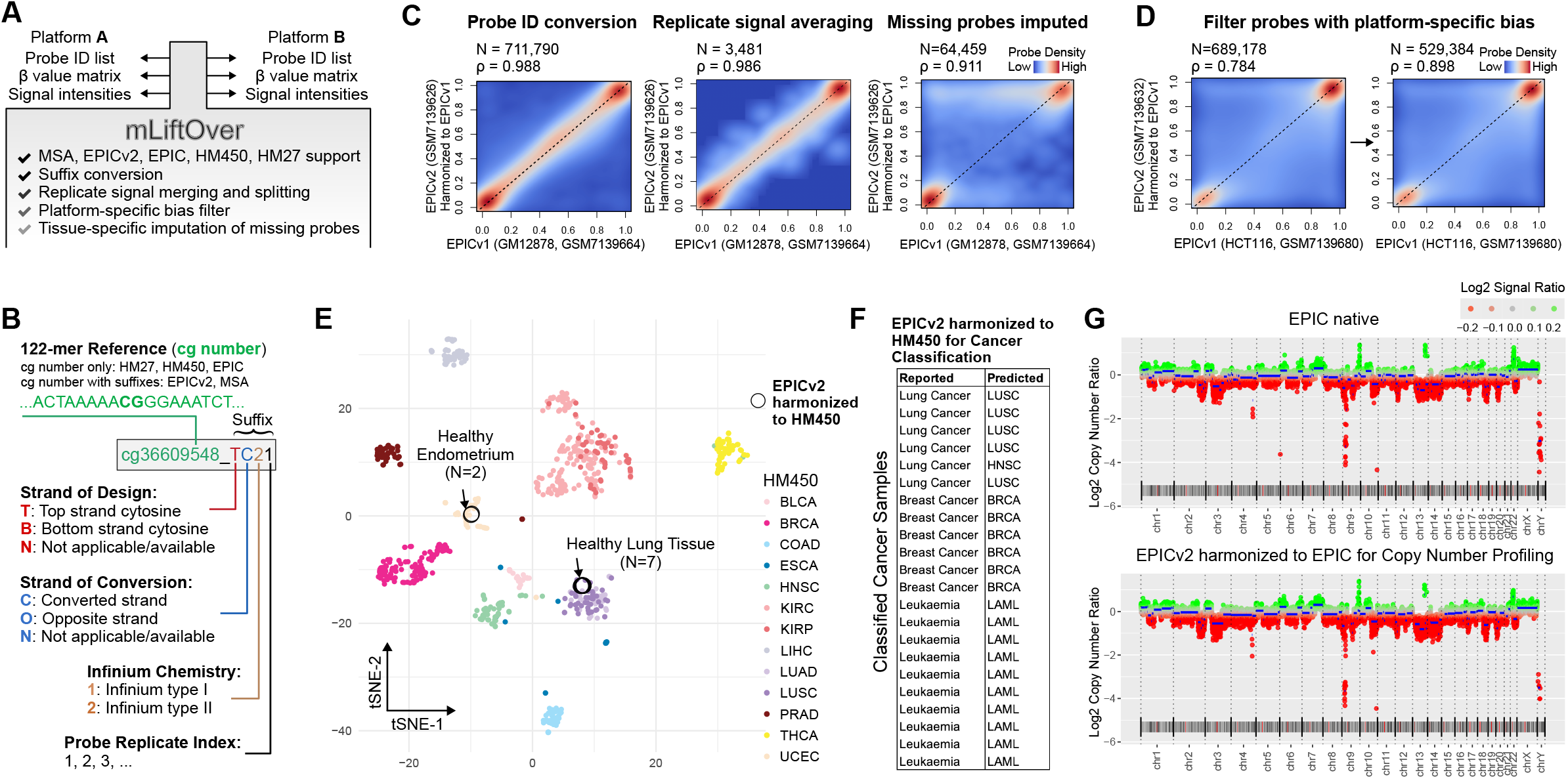
*mLiftOver* harmonizes Infinium DNA methylation BeadChip data across array platforms. (A) Schematic illustration of the core features and workflow of *mLiftOver* from data input to harmonization output. (B) Depiction of the probe naming convention employed in the EPICv2 and MSA arrays. (C) Evaluation of the *mLiftOver*’s accuracy using the GM12878 cell line data, contrasting measurements from EPICv1 and EPICv2. The panel is divided into three sub-panels, demonstrating 1) direct probe ID translation, 2) signal averaging across replicates, and 3) imputation of missing probe readings (excluding those with methylation level standard deviation >0.08). Spearman’s correlation coefficients are displayed atop each subpanel, with all correlations being significant (p-value <1E-6). D) Removal of platform-specific biases (tested on a pair of HCT116 cell line data that did not participate in the platform-specific bias analysis), p-value <1E-6. (E) Illustrates the integration process of *mLiftOver* for primary healthy tissue data and TCGA tumor-adjacent normal tissue data, showcasing its utility in harmonizing diverse datasets. (F) Demonstrates the application of cancer classification models, initially trained on HM450 data using a random forest framework, to primary tumor datasets harmonized from EPICv2 data through *mLiftOver*. (G) Compares copy number variation profiles obtained from native EPIC data and profiles harmonized from EPICv2 data, showing the consistency of *mLiftOver* in signal data conversion.

## 2 DESCRIPTION

*mLiftOver*, developed in R, is a feature in the SeSAMe package (Zhou *et al*., 2018) and leverages the *ExperimentHub* (Pasolli *et al*., 2017) and the *sesameData* packages to organize empirical data for its operation (**Figure 1A**). This tool can convert various data types: Probe IDs as a string list, DNA methylation levels (beta values) as numerical matrices, and signal intensities as *SeSAMe::SigDF* objects. *mLiftOver* is also capable of translating data to and from new and previous Infinium platforms. The tool generically identifies replicate probes as those sharing the same *cg* number prefix but differing in other design aspects, such as strand specification and Infinium chemistry (**Figure 1B**). When integrating data between platforms with and without these suffixes, *mLiftOver* offers two data aggregation strategies: averaging beta values across replicates or selecting the replicate with the most significant signal detection, informed by detection p-values. The latter method can exclude probes with potential design issues as indicated by the mask column within the *SigDF* object. When converting platforms without replicates to platforms with replicates, the same readings will be assigned to different replicates. *mLiftOver* is compatible with all existing Infinium platforms, including HM27, HM450, EPIC, EPICv2, and MM285. It also facilitates the conversion of raw signals stored as *SigDF* class objects, enabling integrated analyses such as copy number variation studies. Beyond signal conversion based on probe IDs, *mLiftOver* can incorporate empirical benchmarks from analyses where two platforms have profiled identical cell lines to filter platform-specific biases, thus enhancing data translation fidelity.

*mLiftOver* integrates publicly available datasets to facilitate the back-conversion of EPICv2 data to its antecedent platforms, EPIC and HM450. This reverse conversion process involves three steps: translating probe IDs, filtering platform-specific biases, and imputing missing data by mapping the sample using the nearest neighbor approach to samples within our comprehensive DNA methylome repository. By aligning with the closest matching tissue type, *mLiftOver* fills in gaps without relying on methylation levels from other samples in the dataset, thereby enabling single-sample dataset operations. We have conducted extensive analyses on 10,631 EPIC and 10,726 HM450 samples to establish a robust imputation baseline when either EPIC or HM450 is the target platform (**Supplemental Table S1**). This baseline collection of datasets spans 20 and 19 tissue types for HM450 and EPIC datasets, respectively, with blood as a focal tissue due to its prevalence in EWAS studies (**Figure S1A**). Additionally, we calculated the variance of beta values for each CpG site within the target tissue type to gauge the confidence of imputation for probes completely absent from the original array. **Figure S1B** shows the standard deviation distribution by tissue type and assay platforms. These variance metrics are critical as they can serve as filters to eliminate methylation influences stemming from unaccounted variables, such as age. The imputation reference data is housed within the *sesameData* package, accessible via the *sesameDataGet* function. When *mLiftOver* detects missing data, it substitutes these gaps with the median methylation value for the respective tissue type. This tissue type is either deduced algorithmically or specified by the user, thereby ensuring the replaced values align with the most probable biological context.

## 3 RESULTS

To show the performance of *mLiftOver*, we benchmarked the accuracy of converted probe-level methylation readings using the EPIC and EPICv2 data profiling the same cell lines (GM12878, K562, and LNCaP) (Kaur *et al*., 2023). We first compared native EPIC data and converted data from EPICv2, then native EPICv2 data and harmonized data from EPIC, all profiling the same cell line (GM12878 or HCT116) (**Figure 1C**). Conversions in both directions highly correlate with the native measurements from the target platform (Spearman *ρ*=0.988) (**Figure 1C**, first panel, **Figure S1C**). EPIC to EPICv2 conversion yields more probes due to the replicate probes with the same *cg* number prefix in EPICv2. Next, compared to native EPIC data, both replicate probe aggregation methods yielded similarly high measurement accuracy on 3,481 probes with design replicates in EPICv2, with the methylation level averaging method slightly surpassing the detection p-value method (**Figure 1C**, second panel, **Figure S1D**). For EPICv2 to EPIC conversion, we further considered data imputation. The imputed values alone also highly correlated with the native EPIC data (Spearman *ρ*=0.82), albeit lower than in the probe sets of direct probe conversion (**Figure S1E**). The Spearman’s correlation remains at 0.977 for converted measurements and imputed values combined (**Figure S1F**). Filtering out 86,678 probes with higher methylation variation (SD>0.08) in the public datasets reduces the number of imputed readings but increases the overall correlation to greater than 0.9 (**Figure 1C**, third panel). Lastly, we tested the filtering of platform-specific biases (**Figure 1D**). We first examined five experiment pairs on three cell lines (GM12878, K562, and LNCaP). We defined a set of high-confidence mapping as those with delta methylation levels no greater than 0.05 in four experiment pairs (see Data Availability). This yielded a mapping of 542,491 EPICv2 probes with 539,513 EPIC probes. *mLiftOver* then uses this mapping to convert unpaired EPIC and EPICv2 experiments on the HCT116 cell lines grown from different labs (Kaur *et al*., 2023). The conversion with the empirical filter yielded a slightly higher correlation (0.898 vs 0.784) with the native data than without filtering platform-specific bias (**Figure 1D**).

To demonstrate the utility of *mLiftOver* in integrating Infinium data across multiple platforms, we applied it to integrate EPICv2 and HM450 data that profiled primary healthy tissue samples. We downloaded two healthy endometrium tissue methylomes and seven lung tissue methylomes (Noguera□Castells *et al*., 2023). We co-clustered the *mLiftOver*-converted methylomes with HM450 datasets of tumor-adjacent normal tissues from The Cancer Genome Atlas (TCGA). As shown in **Figure 1E**, the EPICv2-originated datasets correctly cluster with the corresponding lung and endometrium tissue samples, respectively. This suggests that *mLiftOver* faithfully maintained the tissue-specific signature and epigenetic identity of these biological samples.

Next, we evaluated whether cancer classification models trained on HM450 data can be used on *mLiftOver*-harmonized methylomes. We downloaded 22 primary cancer methylomes of lung cancer, breast cancer, and leukemia (Noguera□Castells *et al*., 2023) and applied a random forest classifier trained on 33 TCGA cancer types (**Figure 1F**). The HM450-based classifier accurately predicts the cancer types of these methylomes except one, leading to an accuracy of 95%.

Lastly, we tested the functionality of *mLiftOver* in converting signal intensities. Infinium array signal intensities are extensively used in discovering copy number aberrations. We benchmarked this functionality on EPIC and EPICv2 datasets profiling the K562 cell lines, a leukemia cell line associated with a characteristic copy number gain at chromosome 22 and loss of chromosome 9p (Zhou *et al*., 2019). As expected, *mLiftOver* can produce consistent copy number profiles from EPICv1 native and EPICv2-harmonized data, capturing this hallmark structural variation (**Figure 1G**).

Collectively, we demonstrate that *mLiftOver* enabled the integration of recent Infinium data with legacy data and allowed for legacy predictive models to be continuously used on data from updated platforms.

## 4 DISCUSSION

The Infinium DNA methylation BeadChip has evolved significantly since its inception, with each subsequent generation enhancing genome coverage while ensuring backward compatibility with existing datasets and models. This progression is exemplified by the seamless transition from the HumanMethylation450 (HM450) to the EPIC array, where most HM450 probes were incorporated into EPIC. Similarly, EPICv2 retained a substantial proportion of probes from EPIC (83%) and HM450 (81%) to preserve continuity. Nonetheless, challenges persist in predictive modeling or longitudinal studies, where comparative analyses are ideally conducted using the same platform. The introduction of replicate probes and more adaptable probe designs has led to the addition of suffixes to the conventional *cg* numbers, hindering the transfer of data across different platform generations and maximizing the use of historical data and models. In this study, we introduce a user-friendly tool designed to streamline data harmonization across three dimensions: probe names, beta values (methylation levels), and raw signal intensities.

With the introduction of new probes and the removal of others in the newer platform iterations, the necessity for imputing missing probe readings has arisen. Our tool, *mLiftOver*, addresses this need by harnessing publicly available data, primarily focusing on tissue-specific differences, which have been identified as principal influencers of DNA methylation patterns in various studies, including our own (Ding *et al*., 2023; Zhou *et al*., 2022). However, we acknowledge that other factors, such as age, sex, cellular malignancy, and mitotic history, have not been incorporated into our model. Moreover, our approach only supports target platforms with enough available data, and tissues with uncharacterized methylomes are absent from our reference database, posing a potential limitation. One possible solution is to utilize the methylation correlation structure, for instance, inferring methylation levels in genomic proximity, to aid in imputing missing data. This approach could exploit co-methylated regions identified in comprehensive genome-wide methylome analyses (Sofer *et al*., 2013). The feasibility of imputation could inform the design of future Infinium arrays. It is important to note that while DNA methylation levels can be imputed, the imputation of signal intensities for absent probes is not yet supported, potentially impacting the analysis of copy number alterations in converted versus native datasets. Nonetheless, *mLiftOver* addresses the problem of probes missing completely between array platforms by utilizing a large database of publicly available DNA methylation array data across multiple tissues and leveraging the variability in methylation levels to assess the imputation accuracy. Our imputation solution for entirely missing probe values can be helpful for predictive models requiring specific probe values, where the alternative would be a missing value. In sum, *mLiftOver* provides user-friendly functionality for projects seeking to analyze DNA methylation data using different versions of Infinium arrays.

