## Supplemental Figure S1 for "mLiftOver: Harmonizing Data Across Infinium DNA Methylation Platforms"

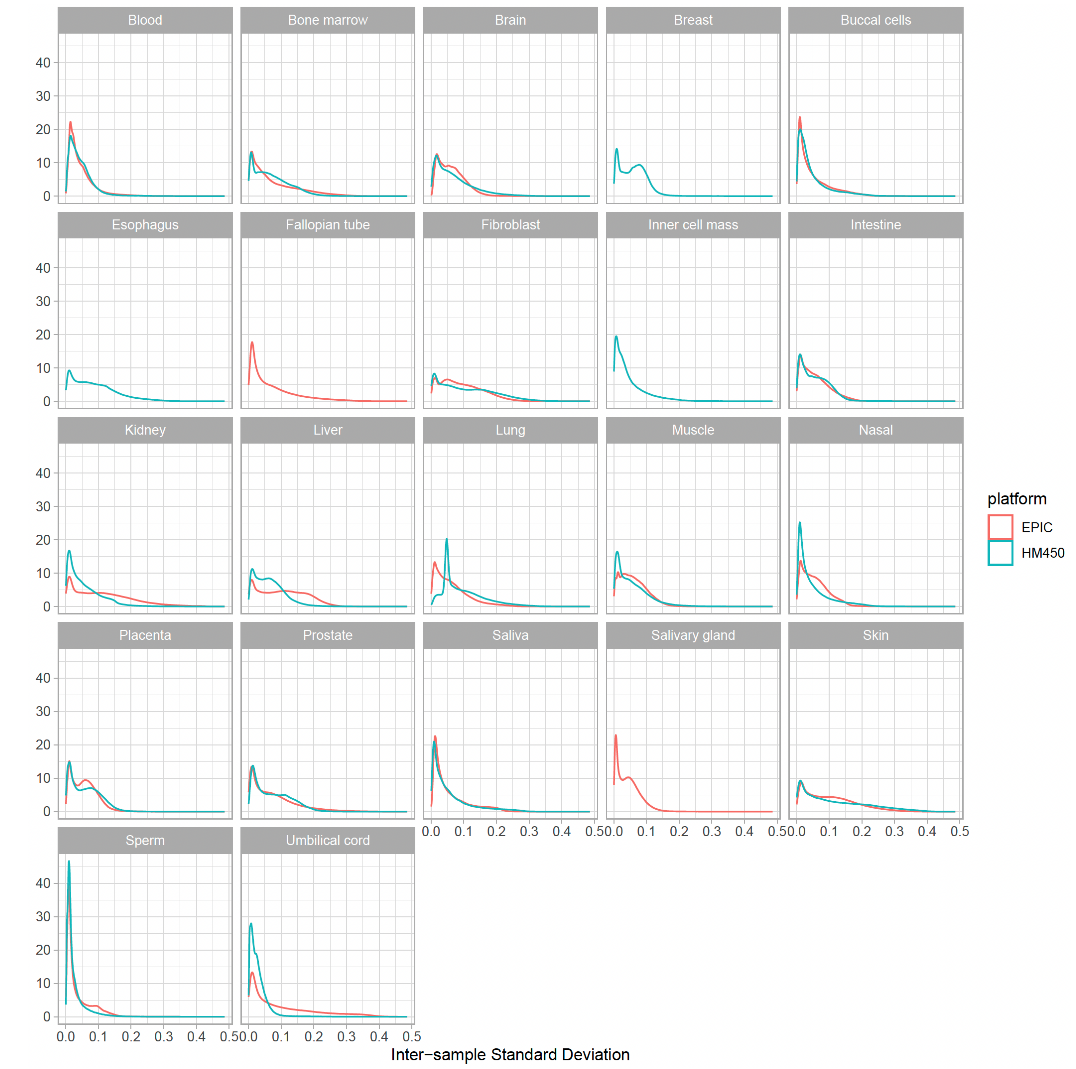


B

| Tissue Type | HM450 | EPIC |
|---|---|---|
| Blood | 2000 | 2000 |
| Bone | 29 | 0 |
| Bone marrow | 281 | 72 |
| Brain | 2000 | 1967 |
| Breast | 288 | 0 |
| Buccal cells | 705 | 446 |
| Esophagus | 59 | 0 |
| Fallopian tube | 0 | 23 |
| Fibroblast | 333 | 327 |
| Inner cell mass | 55 | 0 |
| Intestine | 426 | 1246 |
| Kidney | 119 | 80 |
| Liver | 214 | 232 |
| Lung | 400 | 816 |
| Muscle | 88 | 518 |
| Nasal | 264 | 293 |
| Placenta | 1292 | 1514 |
| Prostate | 67 | 45 |
| Saliva | 509 | 305 |
| Salivary gland | 28 | 103 |
| Skin | 174 | 438 |
| Sperm | 339 | 178 |
| Stomach | 33 | 0 |
| Umbilical cord | 1023 | 28 |

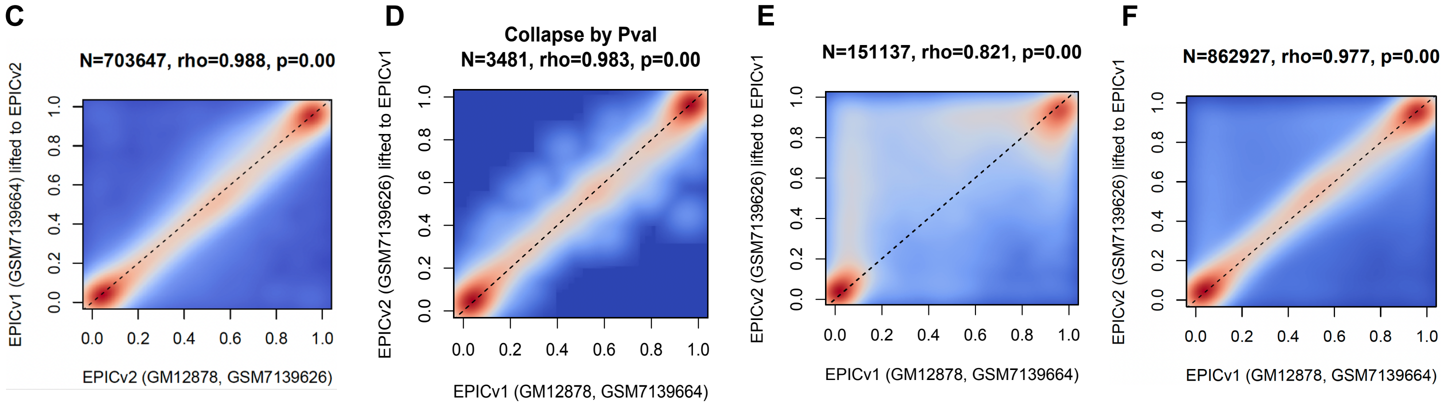


**Supplemental Figure S1. mLiftOver harmonizes Infinium DNA methylation BeadChip data across array platforms.** (A) Methylation level standard deviation in datasets stratified by tissue type and Infinium platforms (HM450 and EPIC). (B) Count of datasets (samples) used to derive baseline methylation imputation reference. (C) Evaluation of mLiftOver accuracy in translating cell line data from EPICv1 to EPICv2 (Y-axis), benchmarked against native EPICv2 data (X-axis). (D) Aggregation of replicate probes by minimal detection P-value. (E) imputation of missing probe readings without exclusion by methylation level standard deviation. (F) Overall accuracy, including both directly translated and imputed probe readings.
